# LazyNet: Interpretable ODE Modeling of Sparse CRISPR Single-Cell Screens Reveals New Biological Insights

**DOI:** 10.1101/2025.06.11.658833

**Authors:** Ziyue Yi

## Abstract

We present LazyNet, a compact one-step neural-ODE model for single-cell CRISPR A/I that operates directly on two-snapshot (“pre → post”) measurements and yields parameters with clear mechanistic meaning. The core log–linear–exp residual block exactly represents multiplicative effects, so synergistic multi-locus responses appear as explicit components rather than opaque composites. On a 53k-cell × 18k-gene neuronal Perturb-seq matrix, a three-replica LazyNet ensemble trained under a matched 1-hour budget achieved strong threshold-free ranking and competitive error (genome-wide r≈0.67) while running on CPUs, compared with transformer and state-space baselines trained on a single V100 with the same time cap. A T-cell screen included only for generalization showed the same ranking advantage under the identical evaluation pipeline. Beyond prediction, LazyNet exposes directed, local elasticities; averaging Jacobians across replicas produces a consensus interaction matrix from which compact subgraphs are extracted and evaluated at the module level. The resulting networks show coherent enrichment against authoritative resources (e.g., large-scale co-expression and curated functional associations) and concordance with orthogonal GPX4-knockout proteomes, recovering known ferroptosis regulators and nominating testable links in a lysosomal–mitochondrial–immune module.

## 1. Introduction

Single-cell RNA sequencing (scRNA-seq) has transformed transcriptomics by revealing the rich cell-to-cell heterogeneity masked in bulk averages, uncovering rare sub-populations, transitional states, and lineage hierarchies across tissues and species [1-2]. Yet each cell is still frozen in time —captured under one environmental condition and at a single instant—so most temporal trajectories and stimulus-response behaviours remain invisible. Systematically probing these dimensions demands direct perturbations: transcriptional read-outs following targeted genetic edits illuminate how regulatory circuits maintain identity, pinpoint druggable control nodes, and expose synergistic gene pairs that can amplify combination therapies or guide precise cell-engineering strategies [18-31]. CRISPR activation or inhibition screens joined with scRNA-seq (“Perturb-seq”) meet this need by bar-coding guide RNAs so that every sequenced cell records its own perturbation history, enabling thousands of knock-outs, knock-downs, or activations to be multiplexed in one experiment [3-4, 32]. The resulting high-dimensional maps of genotype-to-phenotype relationships have already sharpened target validation pipelines, suggested rescue strategies for disease mutations, and provided training data for machine-learning models that aspire to predict unseen edits. These advances have converged in ambitious blueprints for an in-silico “AI virtual cell” (AIVC) capable of simulating cellular behaviour across modalities, contexts, and perturbations [5].

Several recent frameworks aim to predict perturbation responses from single-cell data, including transformer-based models (e.g., scGPT), large foundation models (e.g., CellFM), and graph-based approaches (e.g., GEARS) [6–8]. While powerful, these pipelines typically require multi-GPU hardware, substantial storage, and/or curated pathway priors, and their latent representations are difficult to map onto concrete biochemical rate laws — complicating mechanistic interpretation in routine lab settings. Moreover, recent analyses indicate that deep models do not universally outperform simpler baselines for gene-perturbation prediction, underscoring the need for methods that are both accurate and practical [49]. Neural-ODE approaches offer improved mechanistic fidelity but often assume fixed functional forms or entail numerically intensive integration over high-dimensional states, which limits tractability at transcriptome scale [12–16,50–51].

LazyNet addresses this gap with a neural ODE that embeds a log–linear–exp residual block inside a single explicit-Euler step. Working in log space compresses multiplicative gene–gene effects into a sparse, directly interpretable rate matrix, avoiding attention-style quadratic costs and heavy ODE solvers. Crucially, the model is designed for CPU-level efficiency (no GPU required), operates directly on count matrices without external pathway priors, and matches the common two-snapshot CRISPR A/I design by treating the pre- and post-perturbation profiles as a finite-difference sample of the underlying dynamics. In this study we show that LazyNet attains state-of-the-art predictive accuracy on a large Perturb-seq matrix while training in hours on a single 24-core CPU node, and that its elasticity-based analysis yields compact, biologically coherent subnetworks. Applied to ferroptosis, the framework recovers established regulators and nominates a lysosomal – mitochondrial– immune module for experimental follow-up, illustrating how CPU-tractable, interpretable modeling can turn routine perturbation screens into causal hypotheses at transcriptome scale.

## 2. Materials and Methods

### 2.1 LazyNet Architecture

From Figure1, the full gene-expression vector (INPUT) is propagated through a two-stage transformation composed of a logarithmic layer (LOG) followed by an exponential layer (EXP). These twin layers realise multiplicative biology-inspired interactions in log space while retaining computational simplicity. The transformed signal is added back to the original input via a residual skip connection (black bar and “+”), producing an updated vector of identical dimensionality (OUTPUT). Stacking such blocks corresponds to explicit Euler integration of an underlying ordinary differential equation, allowing LazyNet to model gene-regulatory dynamics within a ResNet framework while continuously adjusting weights during training [17].

**Figure 1.**
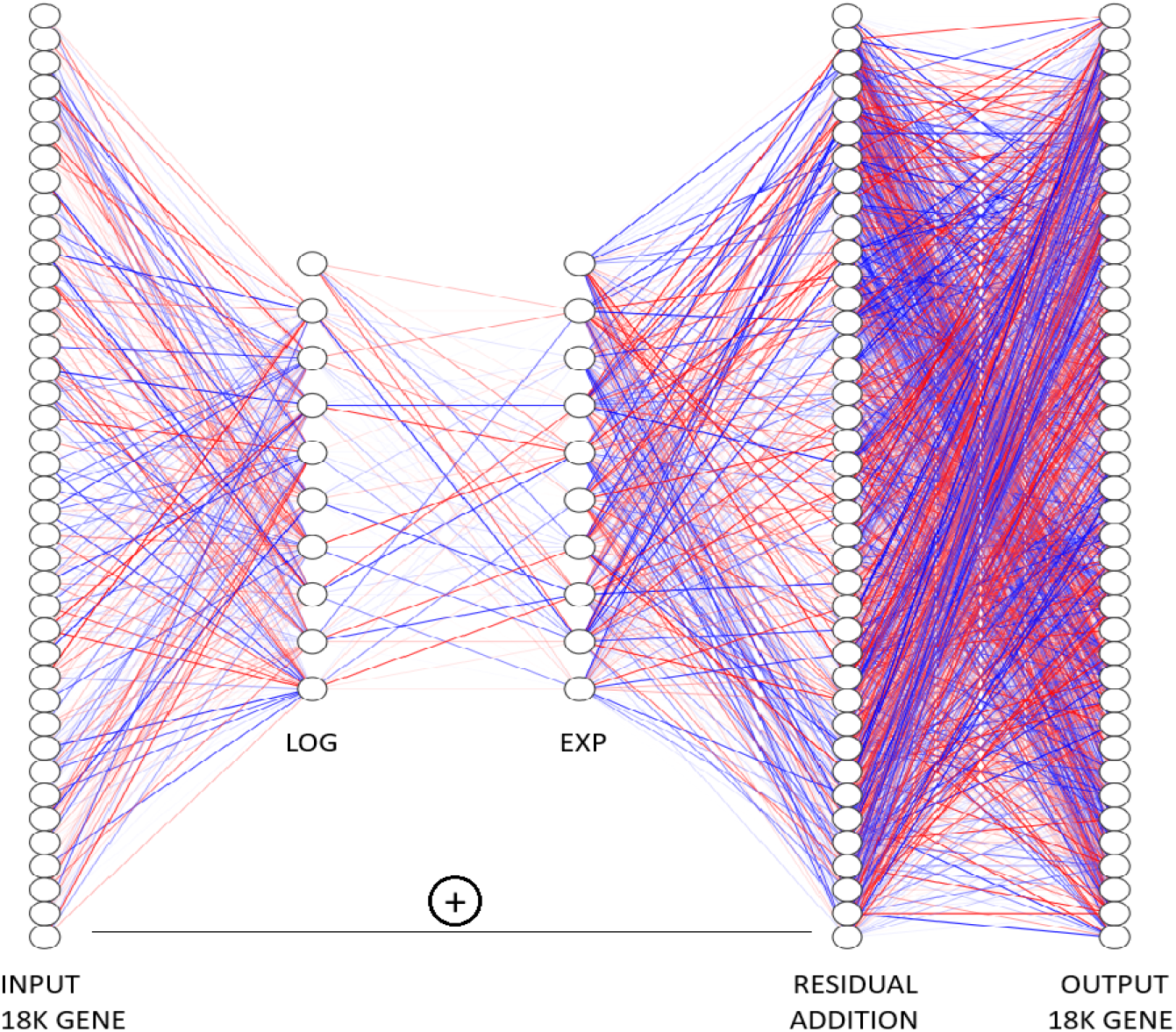
The architecture of LazyNet. All log and exp operations are elementwise; inputs are library-size normalized and shifted by a small pseudocount ε to ensure positivity.

LazyNet has already been applied successfully to both synthetic ODE benchmarks and real-world dynamical systems. Its core idea is to embed a log → linear → exp residual block inside explicit Euler approximation, so that every weight acts as a direct ODE rate constant and the network learns interpretable dynamics with minimal parameters.

An ODE of the form can be discretised with forward Euler:

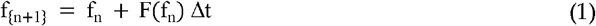

LazyNet represents the rate law F with a single log–linear–exp (LLE) block:

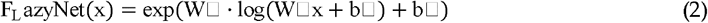

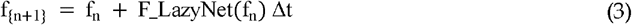

Here x ∈ ℝ +⍰; W_1_, W_2_ are learnable weight matrices and b_1_, b_2_ are biases. Taking the logarithm converts multiplicative gene–gene interactions into additions, allowing a single neuron to encode an entire monomial exactly, while the outer exponential restores the original scale. More details and examples can be seen in Supplement S4.

During training, each observed state fn is provided as input, the network predicts fn+1, and the prediction is compared with ground truth. With sufficiently varied samples V and adequate temporal coverage T, the learned mapping approaches the true governing equations:

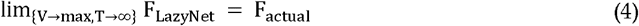

Because the ODE is Markovian, every update depends only on the current state, not on the full history; LazyNet therefore learns the system dynamics efficiently and with minimal parameter count.

In a CRISPR-A/I screen, the before-and-after expression profiles can be treated as Euler approximation of the underlying ODE—thereby sampling the true regulatory function Factual directly.

### 2.2 Theoretical Rationale for Efficiency

Most biochemical rate laws can be written as a finite sum of monomials of the form c · x1α1 … x ⍰αd. A representation that reproduces each monomial exactly captures the full reaction space. LazyNet achieves this with a single log–linear–exp (LLE) block(2), where x ∈ ℝ+⍰, W ∈ ℝK×d, and b ∈ ℝK. Choosing one row of W equal to the integer exponent vector α and the corresponding bias b = log□c reproduces the monomial exactly. Hence an entire K-term rate law needs only K(d□+□1) trainable scalars.

By contrast, a degree-N Taylor polynomial on the same positive domain requires ∼ log□1εd parameters to reach uniform error ε. Thus the parameter ratio grows super-polynomials with both dimension d and desired accuracy. Fewer parameters translate to fewer multiply–add operations per mini-batch and a smaller memory footprint, directly accelerating training.

Statistically, concentrating capacity onto a small set of mechanistically meaningful weights improves the effective signal-to-noise ratio: each observation updates compact exponent and coefficient parameters rather than being diffused across a large dictionary. Empirically, this manifests as faster convergence at larger learning rates, better small-sample generalization, and more stable optimization under residual connections.

Importantly, the multiplicative form encodes synergistic (higher-order, multi-locus) interactions as single monomials rather than as emergent products of many additive units. This is not only an efficiency gain; it directly addresses a substantive modeling need by making synergic effects first-class, precisely parameterized objects that can be learned robustly from limited data.

Transformer-based baselines such as scGPT and state-space models such as RetNet avoid the explicit coefficient blow-up of polynomial bases but incur other costs [6–7]. Attention layers scale as O(L^2^) in sequence length L, while state-space kernels move toward linear time yet introduce millions of additional parameters. Neither family encodes a monomial in closed form; multiplicative gene–gene terms are learned implicitly via gradient descent, typically demanding more data and iterations. In contrast, the LLE map captures multiplicative structure exactly with a forward pass that is linear in d (and in the retained term count K), yielding a favorable accuracy– efficiency–interpretability trade-off.

For AIVC analyses specifically, prior work indicates that LLM-based frameworks (e.g., scGPT, CellFM, GEARS) are not tailored to the dynamics of multi-perturbation, high-order synergy and often struggle to represent such effects faithfully [49]. Neural-ODE approaches can be accurate but usually require a pre-specified differential form—e.g., fixed symbolic templates or universal Hill-type non-linearities [12–16, 50–51]—which constrains adaptability and can produce models that are numerically correct yet mechanistically misspecified. LazyNet retains an ODE viewpoint without fixing the functional form: placing the LLE block inside an explicit-Euler residual update allows approximation of a broad class of interaction terms, so synergy is captured through learned exponents and weights rather than hand-engineered equations, improving both flexibility and mechanistic interpretability.

### 2.3 scRNA Data Preparation and Gene Selection

#### Neuronal Perturb-seq (GSE152988)

We analyze GSE152988, a CRISPR activation/interference screen in 53,495 human iPSC-derived neurons (67,077 genes including non-coding RNAs) with two expression snapshots per cell (a pre-state “time-0” and a post-perturbation state) [33]. With no intermediate measurements, we treat the post-state as a single explicit update from the baseline, matching the minimal ODE setting used throughout this work. No external dynamics or pathway priors were introduced.

To construct “time-0” without leakage, we do not edit individual cells. Instead, we first compute an averaged baseline cell from library-size–normalized non-targeting controls (per batch/stratum as applicable). For each perturbation targeting gene g, the time-0 input is this averaged baseline vector with a single edit at g to encode the intervention: for CRISPRi/KO we set g to a small floor; for CRISPRa/OE we set g to the observed post-perturbation level of g (estimated across the perturbed cells), while all other genes remain at control baselines. Because only the scalar for the targeted gene is updated—and never any multi-gene pattern or any individual test cell—this captures the intended one-gene intervention while preventing multigene peeking into the post-state.

After time-0 construction, we restrict to 18,152 Ensembl GRCh38 protein-coding genes for modeling and evaluation. Cells are grouped by (target gene × gRNA) (and by batch when available), and whole groups are assigned to train/validation/test at 80%/10%/10% to avoid guide- or batch-specific leakage across splits. On the working expression scale used for modeling, the held-out neuronal iPSC test targets have mean 0.6082 and standard deviation 3.212 (min 0, max 1203) as the data landscape.

Because only two snapshots are available, absolute rates are not identifiable; the product of step size and elasticity is what can be learned. Accordingly, downstream evaluation emphasizes one-step prediction errors and relative signals (e.g., rankings) that are invariant to monotone rescaling of the latent time step. When dense time series are available, a multi-step fit resolves Δt and improves identifiability (see Discussion and [14-17]).

#### T-cell guide-capture screen (GSE190604)

We further use a large human T-cell CRISPR guide-capture dataset (10x Genomics RNA with per-cell guide calls) comprising 36,601 genes × 103,805 cells [54]. Expression matrices come from the “Gene Expression” count matrix and guide calls from the corresponding CellRanger export. Minimal QC is applied (genes detected in ≥3 cells; cells with ≥200 detected genes) to preserve a blind and reproducible comparison.

Each barcode is joined to its guide-call summary. Baseline cells (0/unknown guides) are used only to estimate time-0. Singlets (exactly one guide) form the training set; multiplets (≥2 guides) form the test set. After matching and filters, the split yields ∼61k training cells and 28,707 test cells. Counts are library-size normalized with a small pseudocount and log-transformed. We select the top 25% highly variable genes (HVGs) across cells (∼one quarter of 36,601 genes) and carry this fixed HVG panel into all downstream modeling and evaluation. On this normalized log scale, the held-out test targets have mean ≈ 0.19 and SD ≈ 2.30 (min 0, max 2620), which we report to orient error magnitudes in later sections.

### 2.4 Baseline Models

We compared LazyNet to two public baselines chosen to span the principal inductive biases for single-cell perturbation prediction without curated priors: a transformer (scGPT) and a state-space model (RetNet/CellFM) [6–8]. These baselines were selected because they (i) have maintained, open implementations with documented end-to-end pipelines; (ii) do not require curated cell-type labels or pathway priors (e.g., GEARS); (iii) are compatible with two-snapshot Perturb-seq inputs; and (iv) have prior use on closely related single-cell prediction tasks.

Together, scGPT and RetNet/CellFM provide architectural diversity—attention vs. state-space dynamics—yielding an appropriate, apples-to-apples contrast with LazyNet under a common data regime. In this pairing, scGPT serves as the widely adopted attention-based baseline, while RetNet/CellFM offers a competitive linear-time alternative within the state-space family.

### 2.5 Training Protocol

LazyNet is trained on our Intel Xeon CPU server Intel Xeon Gold 6126, 24 cores, 768 GB RAM; no accelerators. The main runs used a Huber loss (δ = 0.1) for robustness to outliers and a three-replica LazyNet ensemble (independent seeds; arithmetic averaging at inference). Complete training hyperparameters, exact command lines, and seeds are provided in the replication package.

In the ablations, the model depth is the most sensitive knob: increasing the log-exp pair more than one led to numerical instabilities (NaNs) or higher error, consistent with overfitting in a setting where a one-step ODE is already a low-complexity mechanism and LazyNet is highly expressive. Width is less sensitive, but a moderate range (several hundred units) performs best; widths that are too small underfit, while very wide models tend to overfit or fail to converge usefully within the time budget. The additive residual path is essential for ODE fitting—removing it substantially degrades accuracy or prevents stable training. We also varied the loss family (Huber/L1/L2 and δ), mini-batch size, and integration step length. Within the ∼1 h per-replica budget (3 replicas), batch size and step length had negligible effect on test metrics, whereas expanding LazyNet beyond moderate capacity often undertrained within the time cap, yielding lower accuracy despite more parameters.

scGPT and RetNet/CellFM are trained on a single NVIDIA V100 GPU with three CPU cores allocated. To match LazyNet’s elapsed-time constraint, each baseline is capped at 1 hour of wall-clock time per dataset, inclusive of data loading and evaluation. We rely on the authors’ documented defaults where applicable, enable mixed precision and gradient accumulation as needed to respect memory limits, and do not introduce curated priors or external labels. The last checkpoint produced within the time cap is evaluated on the common test split.

Regression fidelity is assessed with RMSE, MAE, and Pearson’s r on the modeling gene panel defined in § 2.3. Threshold-free ranking quality is summarized by ROC-AUC and PR-AUC computed directly from continuous predictions. When reported, F1 (micro and macro) uses a single per-gene operating point learned once on the training data via a fixed quantile rule and held fixed for evaluation. All models use identical train/validation/test partitions, deterministic data loaders, fixed random seeds, and the same evaluation scripts.

### 2.6 Network Inference

To produce a directed, dynamical snapshot for network analysis, we averaged ten independently trained LazyNet checkpoints into a single ensemble and evaluated it on the full single-cell matrix (train + validation + test). From this ensemble we computed a Jacobian of absolute elasticities at a common baseline expression vector, restricted to the original 18,152 Ensembl protein-coding genes [34]. For each literature-anchored seed gene (seven total; Supplement 1A), we ranked both downstream (row) and upstream (column) elasticities, retained neighbors with |∂x⍰/∂x⍰| ≥ 0.001 that also exceeded either the 95th or 99th percentile of that seed’s ranked list, and then kept the top-N survivors. Because a single-baseline Jacobian yields few candidates per seed, we set the Benjamini–Hochberg threshold to q = 1 (effectively no FDR filtering) and report the filtered ranked sets. Evaluation proceeds in two parts: first, whether each seed’s neighborhood recovers interactions reported in the literature; second, cross-seed recall, i.e., how often other seed genes appear among a given seed’s ranked neighbors.

The goal is to recover directed, dynamical relationships consistent with the ODE-based formulation—edges that reflect functional influence under perturbation, not merely correlation. For this reason, GENIE3/GRNBoost/SCENIC-style pipelines are not used as head-to-head comparators here. They operate as subgraph generators that prioritize local correlation edges and typically require a manual regulator prior (e.g., TF lists) to seed the search [39,41, 43]. Their outputs are commonly undirected and tuned to rank local associations, rather than to assemble a globally coherent dynamic network. They also lack the external authority of curated/experimental resources; when a comparator is needed, databases such as STRING and ARCHS4 — grounded in empirical evidence — are more appropriate reference points than GENIE3-family predictions.

Even with authoritative databases, edge-level comparisons remain problematic: curated graphs are incomplete, context-dependent, and unevenly sampled, and the true functional wiring can vary substantially even within the same cell type and omic snapshot. We therefore evaluate at the module level, where the signal is more robust: given a set of curated modules (pathways, TF regulons), we quantify module activity/enrichment induced by the inferred network (or by model-predicted responses) and test whether the recovered activity patterns align with known biology. This enrichment-based evaluation emphasizes coherent functional programs over individual edges, respects the dynamical and directed nature of our model, and provides a fairer, more stable basis for comparison across methods and references.

### 2.7 ARCHS4 co-expression validation

We annotate each directed edge (i→j) with the ARCHS4 Pearson correlation r(i,j). To control for confounding, enrichment is computed against a matched null: for each edge we sample matched pairs (i′→j′) from strata that jointly bin genes by (i) control-cell detection/mean expression and (ii) out-/in-degree in the LazyNet elasticity graph (≈400 matched pairs per edge; 9,860 edges; 3,943,792 null pairs). We report fractions exceeding |r| ≥ τ and the corresponding fold enrichment (Observed/Expected). For sign-aware checks, we multiply r by the LazyNet edge sign; because ARCHS4 is undirected/heterogeneous, we emphasize |r| results and treat sign panels as exploratory.

### 2.8 STRING interaction concordance

STRING functional associations (human, version 12.0) were converted from protein identifiers to HGNC symbols with the official mapping file [36]. Isoform-specific and unmapped entries were removed. 32*4 LazyNet sub-graph edges were queried for exact symbol matches, and statistical significance of the overlap was assessed with a one-tailed hypergeometric test that treats the total number of possible unordered human gene pairs as the population size. Elasticity scores and STRING combined scores were each dichotomized at the 75th percentile to classify edges as “both-high”, “STRING-only” or “LazyNet-only”. Degree-preserving Maslov–Sneppen rewiring (1000 iterations) provided a null distribution for network-density comparisons.

### 2.9 Proteomics Validation

Pulsed-SILAC proteomes were re-analysed to evaluate 32*4 LazyNet sub-graph at the protein layer. For the tamoxifen-inducible GPX4-knockout time-course in mouse fibroblasts (PRIDE PXD050979), proteinGroups.txt (PXD050979.txt) was taken as input [47]. Reverse sequences and common contaminants were discarded; gene symbols were extracted, converted to upper-case HGNC format and duplicate entries were averaged, yielding 1026 proteins. Intensities for the light, medium and heavy SILAC channels (nine runs) were treated as independent samples, zeros set to missing and imputed with the MinProb left-shift approach (μ – 1.8 σ; width 0.3 σ) as implemented in Perseus [48]. Values were log_2_-transformed, median-centred per run and proteins quantified in at least three channels with non-zero variance were retained.

For the larger constitutive GPX4-knockout study in Pfa1 mouse embryonic fibroblasts (PRIDE PXD040094) we used the combined_protein.tsv [46]. Gene symbols were cleaned as above and duplicate symbols collapsed. Columns labelled “MaxLFQ Intensity” defined fourteen quantitative channels covering wild-type and knockout states. Missing values were replaced with MinProb and the matrix was log_2_-transformed and median-centered. Proteins present in at least three channels and showing non-zero variance were kept.

For each dataset the cleaned protein matrix was intersected with the RNA-based LazyNet edge list. Gene pairs with at least three paired, finite observations were assigned a Spearman rank correlation coefficient. To control for hub bias a degree-matched null of 10k random edges was generated by stratified sampling across twenty-one degree bins (0-to-20+). Absolute correlation values of LazyNet edges were compared with the null using a two-sided Wilcoxon rank-sum test.

### 2.10 Experiment Layout

As shown in Figure 2, we start from a single pre-processed neuronal iPSC Perturb-seq matrix (library-size normalization followed by a small-pseudocount transform, z=log(x+ε)) and run two parallel analyses.

**Figure 2.**
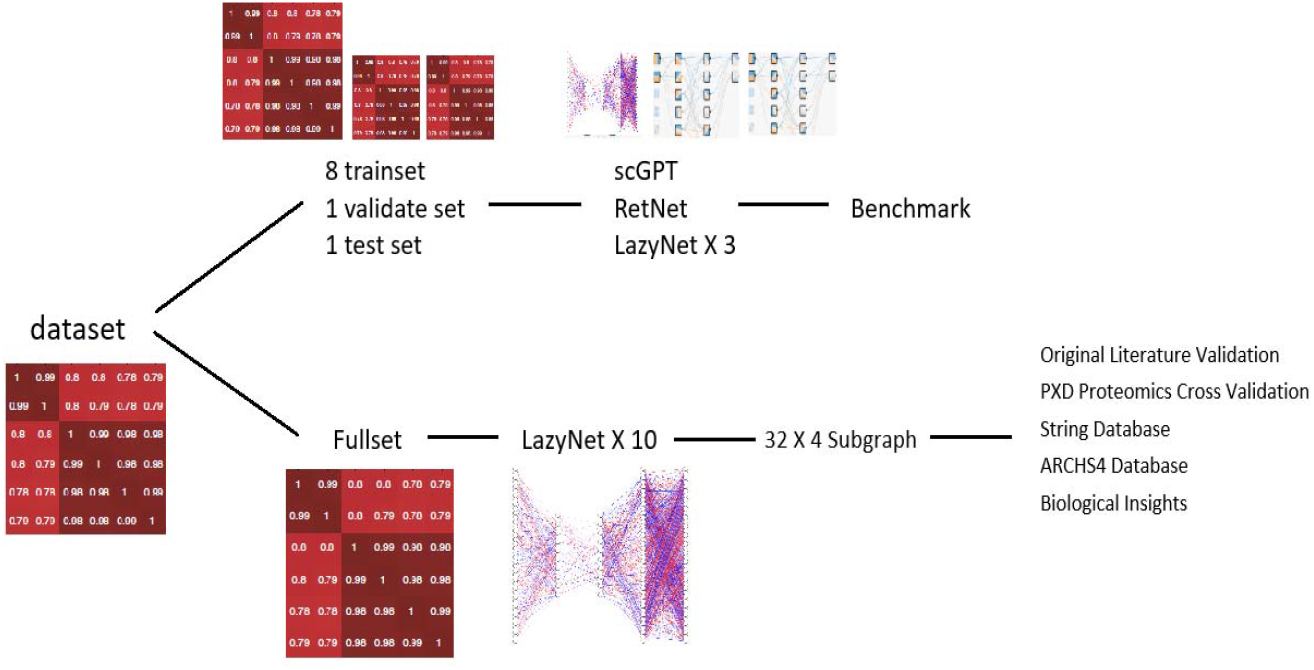
Benchmark-and-discovery workflow.

The benchmark arm performs a one-time stratified split (8:1:1 train:validation:test) and trains scGPT, RetNet, and three LazyNet replicas on the training fold, tunes on validation, and reports performance on the locked test fold (RMSE, MAE, Pearson r; threshold-free AUCs where noted) [6 –8]. A T-cell screen is included only in this benchmark arm (not shown in figure) to test cross-dataset generalization under the identical pipeline; our central goal remains the neuronal dataset.

The discovery arm fits ten independently seeded LazyNet models on the full neuronal matrix (train+validation+test), averages their Jacobians (restricted to 18,152 Ensembl protein-coding genes), and extracts a 32×4 breadth-first subgraph as a consensus network for interpretation.

This subgraph is evaluated against authoritative resources—STRING v12 and ARCHS4 [35–36]— and cross-omic coherence is assessed in two GPX4-knockout SILAC proteomes from mouse embryonic fibroblasts (PRIDE PXD040094, PXD050979) [46–47]. The validated interactions are then mined for mechanistic hypotheses and used to nominate follow-up perturbations.

## 3. Results

### 3.1 Model Benchmark

Table 1 summarizes head-to-head results for a neuronal iPSC Perturb-seq slice for later inference and a T-cell guide-capture slice (top-quartile HVGs) solely for generalization test.

**Table 1.**
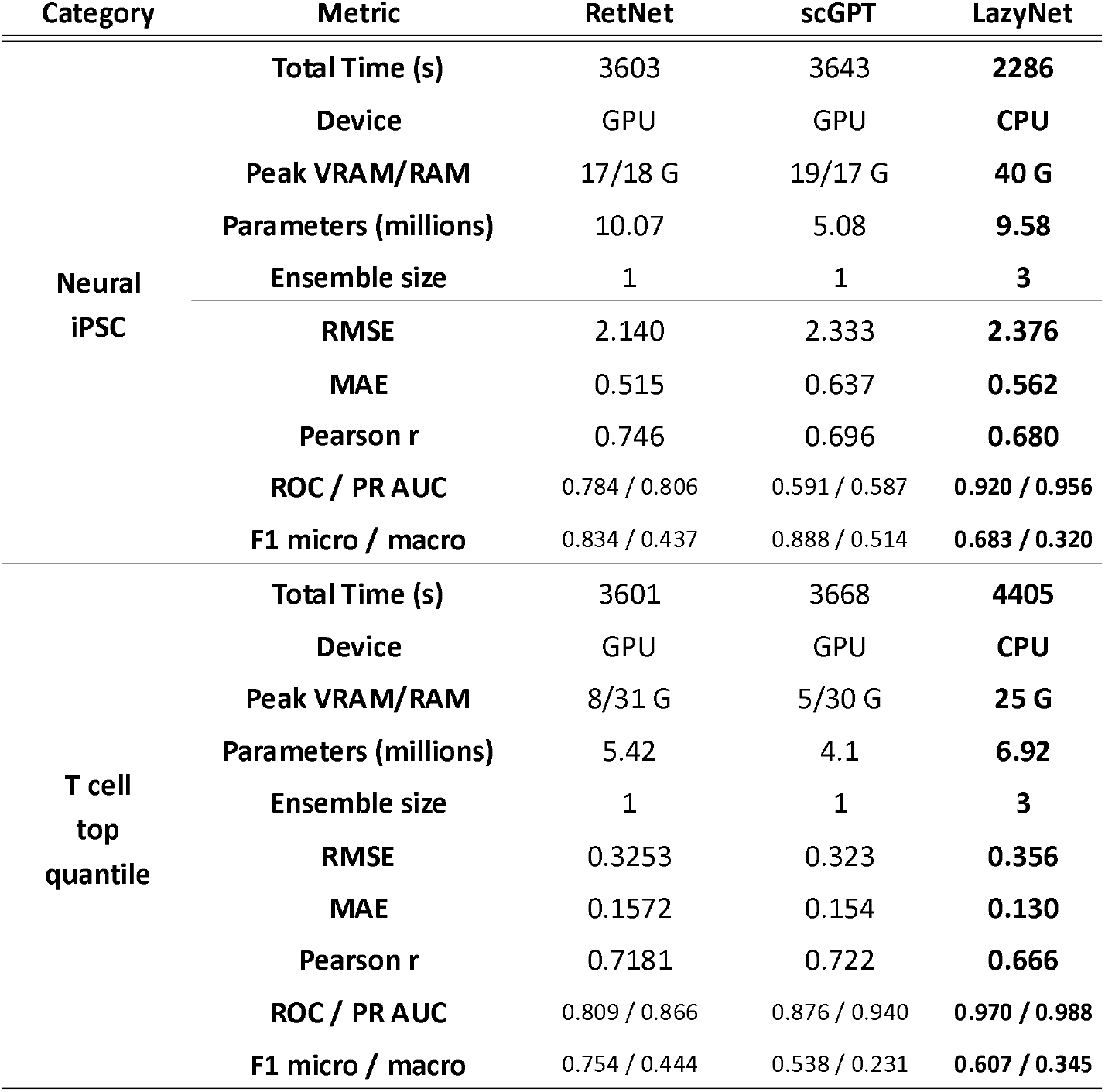
Comparison of training cost and predictive accuracy.

| Category | Metric | RetNet | scGPT | LazyNet |
| --- | --- | --- | --- | --- |
| Neural<br>iPSC | Total Time (s) | 3603 | 3643 | **2286** |
|  | Device | GPU | GPU | **CPU** |
|  | Peak VRAM/RAM | 17/18 G | 19/17 G | **40 G** |
|  | Parameters (millions) | 10.07 | 5.08 | **9.58** |
|  | Ensemble size | 1 | 1 | **3** |
|  | RMSE | 2.140 | 2.333 | **2.376** |
|  | MAE | 0.515 | 0.637 | **0.562** |
|  | Pearson r | 0.746 | 0.696 | **0.680** |
|  | ROC / PR AUC | 0.784 / 0.806 | 0.591 / 0.587 | **0.920 / 0.956** |
|  | F1 micro / macro | 0.834 / 0.437 | 0.888 / 0.514 | **0.683 / 0.320** |
| T cell<br>top<br>quantile | Total Time (s) | 3601 | 3668 | **4405** |
|  | Device | GPU | GPU | **CPU** |
|  | Peak VRAM/RAM | 8/31 G | 5/30 G | **25 G** |
|  | Parameters (millions) | 5.42 | 4.1 | **6.92** |
|  | Ensemble size | 1 | 1 | **3** |
|  | RMSE | 0.3253 | 0.323 | **0.356** |
|  | MAE | 0.1572 | 0.154 | **0.130** |
|  | Pearson r | 0.7181 | 0.722 | **0.666** |
|  | ROC / PR AUC | 0.809 / 0.866 | 0.876 / 0.940 | **0.970 / 0.988** |
|  | F1 micro / macro | 0.754 / 0.444 | 0.538 / 0.231 | **0.607 / 0.345** |

On the neuronal iPSC slice, LazyNet attains the strongest threshold-free ranking quality with the highest ROC-AUC and PR-AUC among the compared methods, while training faster on CPU and using a mid-range parameter count. Its absolute regression errors (RMSE, MAE) and correlation r are slightly worse than the best baseline, and its F1 (micro/macro) is lower at the fixed per-gene operating point—an expected AUC–F1 trade-off when a single threshold is pre-declared per gene.

Together, the iPSC results indicate that LazyNet delivers the best ordering of perturbation effects under CPU constraints, with modestly higher regression error than the top baseline.

On the T-cell HVG slice, LazyNet again leads on ranking metrics with the highest ROC- and PR-AUC, and it achieves the best MAE, while its RMSE and Pearson r are lower than those of the strongest baseline and its F1 sits mid-pack at the fixed threshold (Table 1). This pattern—excellent ranking with conservative amplitudes—matches the expected behavior when per-gene thresholds are learned once on singlets and then applied to a broader test distribution of multiplets.

Interpreting AUC and F1 together clarifies these results. ROC-/PR-AUC capture threshold-free ranking and are insensitive to the exact operating point; F1 fixes a single per-gene threshold learned on training singlets, so it can move counter to AUC when calibration shifts or variance broadens between train and test. In our setting, this explains why LazyNet can lead decisively in AUCs yet not maximize F1 on every slice. All definitions and thresholding rules are held constant across models to keep comparisons fair.

Overall, across both slices and under CPU-only execution with a three-run LazyNet ensemble, the model offers the strongest ranking of perturbation effects and a favorable MAE profile on T-cells, while ceding some ground on RMSE and r to the best baseline on certain metrics. Because LazyNet instantiates a one-step ODE with log–exp residual dynamics, it exposes per-gene elasticities whose ensemble Jacobian yields directed, mechanism-level edges; we leverage this structure immediately in network inference, where the Jacobian is converted into a consensus subgraph and evaluated via module-level enrichment against curated resources.

### 3.2 Network Inference

Ten independently trained LazyNet replicas were averaged into a single ensemble, and Jacobian elasticities were queried around seven ferroptosis seeds (GPX4, PSAP, ACSL4, WIPI2, MTOR, NFU1, SOD2), selection details appear in Supplement S1A. A broader parameter sweep, summarized in Supplement S1B, identified five expansion regimes that were carried forward (Table 2).

**Table 2:**
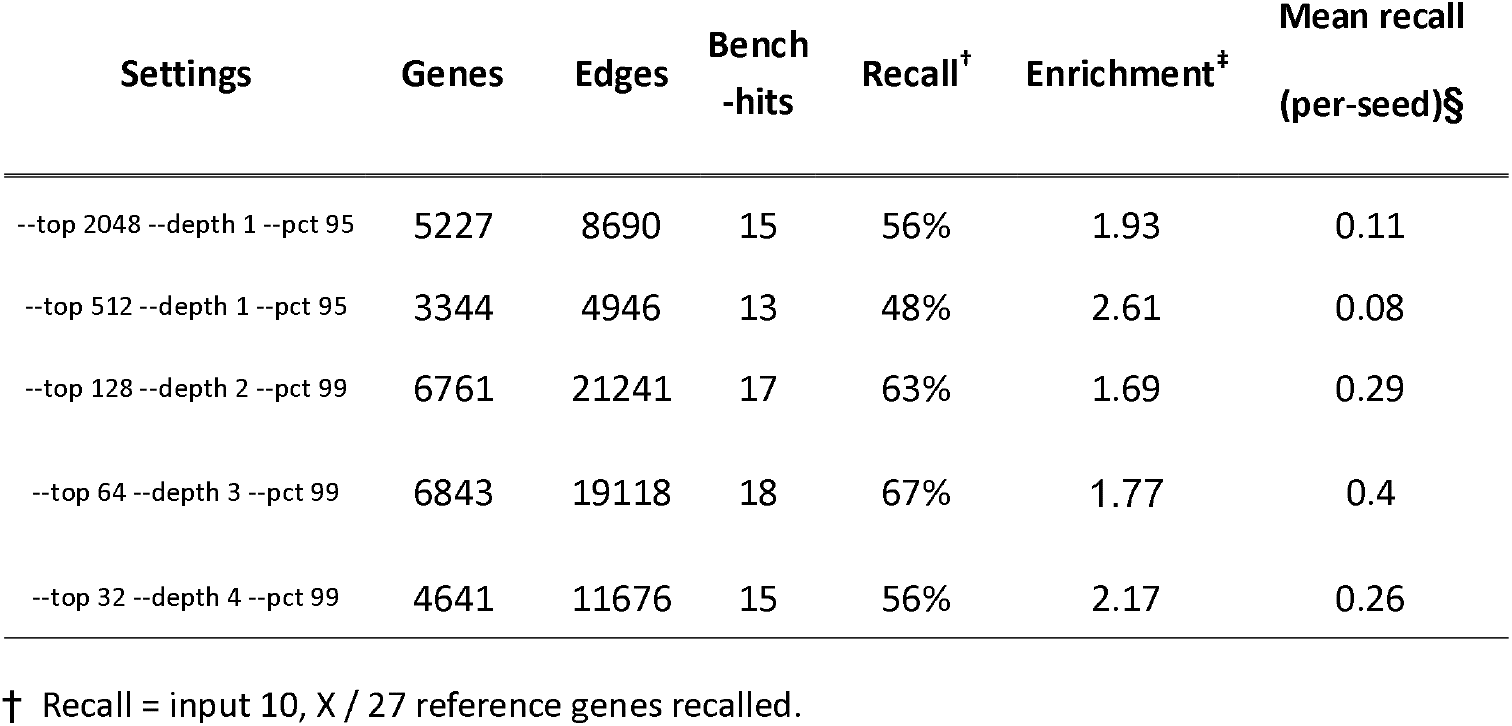
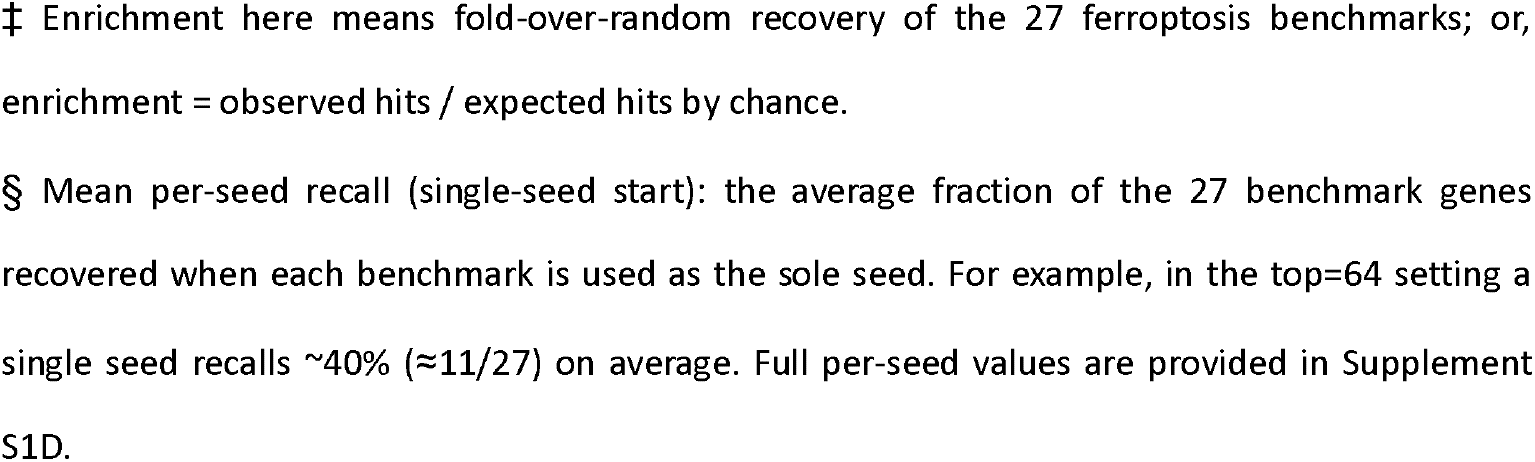
gene inference stats.

| Settings | Genes | Edges | Bench<br>-hits | Recall<sup>†</sup> | Enrichment<sup>‡</sup> | Mean recall<br>(per-seed)<sup>§</sup> |
| --- | --- | --- | --- | --- | --- | --- |
| --top 2048 --depth 1 --pct 95 | 5227 | 8690 | 15 | 56% | 1.93 | 0.11 |
| --top 512 --depth 1 --pct 95 | 3344 | 4946 | 13 | 48% | 2.61 | 0.08 |
| --top 128 --depth 2 --pct 99 | 6761 | 21241 | 17 | 63% | 1.69 | 0.29 |
| --top 64 --depth 3 --pct 99 | 6843 | 19118 | 18 | 67% | 1.77 | 0.4 |
| --top 32 --depth 4 --pct 99 | 4641 | 11676 | 15 | 56% | 2.17 | 0.26 |

<sup>†</sup> Recall = input 10, $X / 27$ reference genes recalled.
‡ Enrichment here means fold-over-random recovery of the 27 ferroptosis benchmarks; or, enrichment = observed hits / expected hits by chance.
§ Mean per-seed recall (single-seed start): the average fraction of the 27 benchmark genes recovered when each benchmark is used as the sole seed. For example, in the top=64 setting a single seed recalls ~40% ( $\approx 11/27$ ) on average. Full per-seed values are provided in Supplement S1D.

In the broadest setting (512×1) each seed ballooned into a neighborhood of ∼3000 genes—GPX4 alone recruited 1811—and yielded a mean per-seed recall of ∼0.08 (≈2–3/27). Constraining the search to 64×3 reduced each subgraph to ∼1000 genes and lifted the mean per-seed recall to ∼0.40 (≈11/27). The most selective regime, 32×4, fused all seven seeds into a single graph of 4641 genes and 11676 edges while preserving a benchmark recall of 15/27 and nudging the hit-rate upward; its data are in Supplement S1C. Because only 27 of 18,152 genes are treated as positives (base rate ≈0.15%), these fractions are numerically small; viewed as enrichment, the 32 ×4 graph recovers 15 regulators versus ∼6.9 expected by chance (∼2.17×).

Evaluate all the gene seeds, per-seed recalls (Supplement S1D) highlight a practical strength: starting from just one ferroptosis gene, LazyNet typically retrieves ∼10 additional benchmarks when run at top = 32, 64, or 128. In other words, minimal prior knowledge suffices to reconstruct much of the ferroptosis module—evidence of the method’s consistency and day-to-day utility. LazyNet recovers 56 % (15/27) of benchmark ferroptosis interactions in < 6 h on a single 24-core CPU node (768 GB RAM, no GPUs). By comparison, GENIE3 recovers ∼33 % of gold-standard E. coli edges after analyzing > 100 microarrays and ∼48 h on 32 cores [39-40]; GRNBoost recalls ∼22 % in a 68 k-cell PBMC atlas [41-42]; and DeepFGRN reaches ∼40 % but requires dense time-series data and multi-GPU training (4*A100, several days) [43]. Thus, LazyNet achieves state-of-the-art recall with orders-of-magnitude fewer time points and far lighter compute while still yielding biologically interpretable subnetworks from a single seed gene.

The 32 × 4 expansion is chosen for all subsequent analyses because it strikes the right balance between biological depth and practical scope. Limiting each hop to 32 of the strongest elasticity neighbors keeps the growth of the graph contained, while four propagation steps are ample to capture multi-step regulatory cascades. The resulting sub-graph (4641 genes and 11676 edges) remains small enough for reliable enrichment statistics, manual curation, and clear visualization, yet large enough to preserve the key ferroptosis signals that LazyNet uncovers. This setting therefore provides a tractable, biologically informative backbone for downstream validation and interpretation.

### 3.3 ARCHS4 Co-expression

Of the 11,676 edges in the 32×4 subgraph, 9,860 (84.4%) mapped to the ARCHS4 pan-human matrix and received a co-expression value (Pearson r) [35]. The r distribution is broadly centered near zero (mean ≈ 0.053; median ≈ 0.044) and closely matches degree-matched random pairs (Supplement S2A).

In an unsigned view, LazyNet edges are far more likely than random to show moderate co-expression: 26.6% of edges meet |r| ≥ 0.20 versus ∼5% of random pairs— an ≈5 × enrichment—indicating that the model links genes that frequently co-vary across thousands of public RNA-seq datasets (Supplement S2B).

Sign-aware analysis refines this picture in Figure 3: the positive, sign-matched tail is enriched (∼1.36 fold), the opposite-sign tail is depleted (∼0.67 fold), and the overall |r| ≥ 0.20 mass is near expectation (∼1.02 fold) once signs are considered (Supplement S2C). This is consistent with the biology: LazyNet elasticities encode local, directional responses in a one-step ODE, whereas ARCHS4 aggregates steady-state, undirected correlations across heterogeneous tissues; inhibitory or context-specific links can therefore appear anti-correlated in the aggregate.

**Figure 3.**
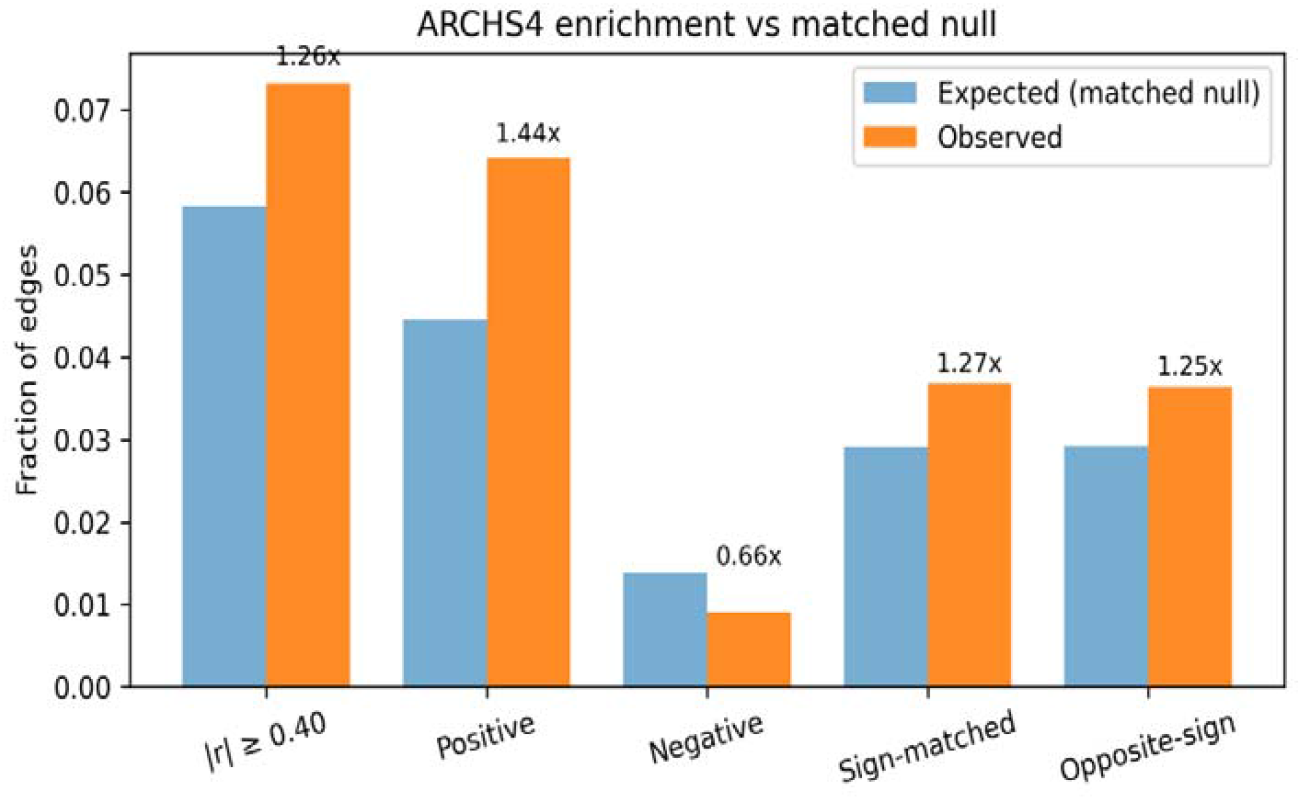
ARCHS4 co-expression enrichment for LazyNet edges. Observed (orange) versus expected (blue) fractions for edges from the 32×4 LazyNet subgraph annotated in ARCHS4 (n = 9,860). To avoid inflation from expression abundance and hubness, expected fractions come from an expression/detection- and degree-matched null (≈3.94M matched pairs). Bars show tail rates at |r| ≥ 0.40; numbers above bars are fold enrichment (Observed/Expected). LazyNet edges are enriched in the overall |r| tail (0.073 vs 0.058; 1.26×) and especially the positive tail (0.064 vs 0.045; 1.44×) with depletion in the negative tail (0.009 vs 0.014; 0.66×). “Sign-matched” multiplies ARCHS4 r by the LazyNet edge sign; “Opposite-sign” reverses it. As expected for an undirected, context-mixed resource, sign-aware fractions are similar (1.27× vs 1.25×).

Taken together, the strong unsigned enrichment and the direction-consistent positive-tail enrichment indicate that a substantial subset of LazyNet edges is corroborated by large-scale expression evidence while respecting expected sign- and context-dependent effects.

### 3.4 STRING Database and Literature Validation

We evaluated the 32 × 4 LazyNet sub-graph against a filtered STRING network containing the 18152 genes detected in our single-cell atlas [36]. Of the 4630 genes retained by LazyNet, STRING supplies 411761 undirected links, giving an observed density of 0.0384. Size-matched random gene sets drawn 1 000 times from the same background average 0.0373 ± 0.0008, placing the sub-graph 1.5 s.d. above expectation (empirical p = 0.072) — evidence for modest but over-connectivity (Supplement S3A).

Edgewise, LazyNet 11 662 elasticity links intersect STRING in 523 cases (Jaccard ≈ 1 × 10^−4^). A one-tailed hypergeometric test confirms this overlap is highly non-random (p = 1.2 × 10^−5^). Although elasticity magnitudes and STRING scores show no global correlation (Pearson r = – 0.071, p = 0.10; Spearman ρ = –0.036), a top-quartile split in each metric separates three informative classes: 27 “both-high” edges, 104 STRING-high only (canonical interactions), and 104 LazyNet-high only—highlighting under-explored relationships for experimental follow-up. Representative literature support for ten both-high edges is provided in Table 3.

**Table 3:**
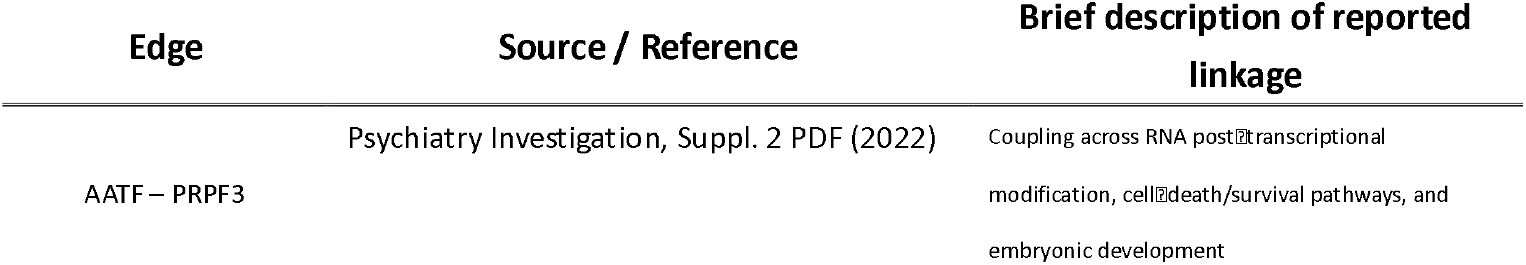
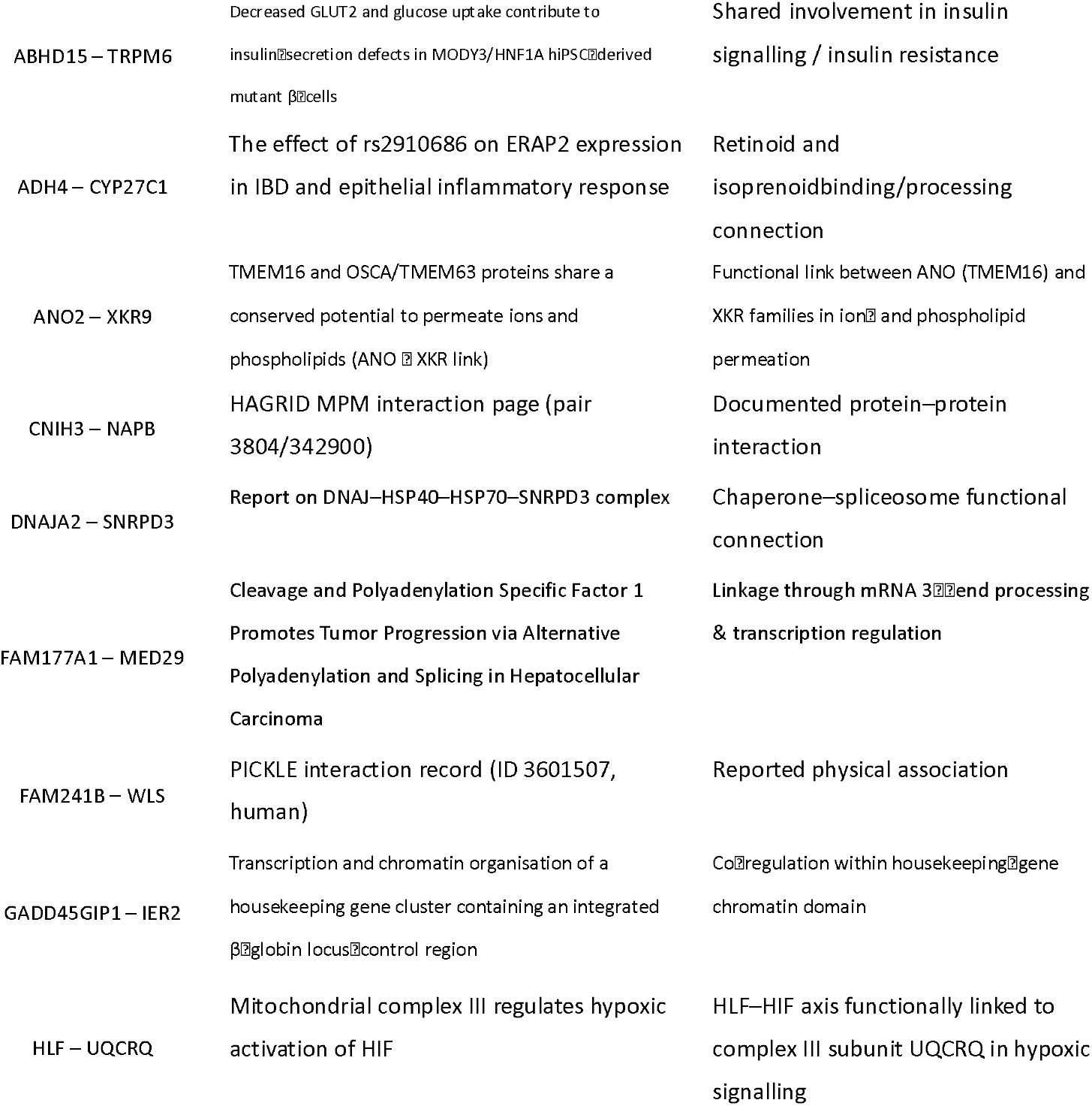
STRING database literature comparison.

Among the findings, the moderate enrichments are entirely acceptable—and even expected— because STRING confidence and LazyNet elasticity quantify different biological facets. STRING aggregates heterogeneous evidence (co-expression, physical binding, literature co-mentions), whereas elasticity captures the transcriptional impact inferred by our ODE model. Perfect concordance would therefore be unrealistic; instead, the modest density excess and significant, yet limited, edge-level overlap demonstrate that LazyNet recovers a meaningful core of known biology while offering a large complement of novel, model-specific interactions.

### 3.5 Proteomics corroborates transcript-derived network structure

Cross-omics robustness was assessed by projecting RNA-derived edges onto two independent mouse fibroblast SILAC proteomes of GPX4 loss-of-function ferroptosis. The inducible GPX4-knockout time course (PRIDE PXD050979) yielded 1,026 quantifiable proteins after consolidation; 47 network edges had sufficient protein coverage (≥ 3 finite values per partner). Their absolute protein co-variation was higher than degree-matched random pairs (median |ρ| = 0.536 vs 0.400; Wilcoxon P = 4.9 × 10^−3^). The larger pulsed-SILAC study of constitutive GPX4 knockout (PRIDE PXD040094) provided 5,516 proteins across 14 channels; of 563 mapped edges, 507 met the coverage criterion and likewise exceeded the null (median |ρ| = 0.367 vs 0.336; Wilcoxon P = 6.2 × 10^−4^). Representative protein–protein scatters are shown in (Figure 4), including XDH–ANXA8 (ρ = 0.92), RPLP2–BAIAP2 (ρ = 0.89), NRDC–ENO3 (ρ = 0.88), and SFPQ– CKAP5 (ρ = 0.98), illustrating coherent proteomic co-variation across channels. Together with the enrichment tests, these results indicate that a substantial subset of LazyNet edges remains concordant at the protein layer, supporting the biological fidelity of the inferred network, despite differences in species and experimental design.

**Figure 4.**
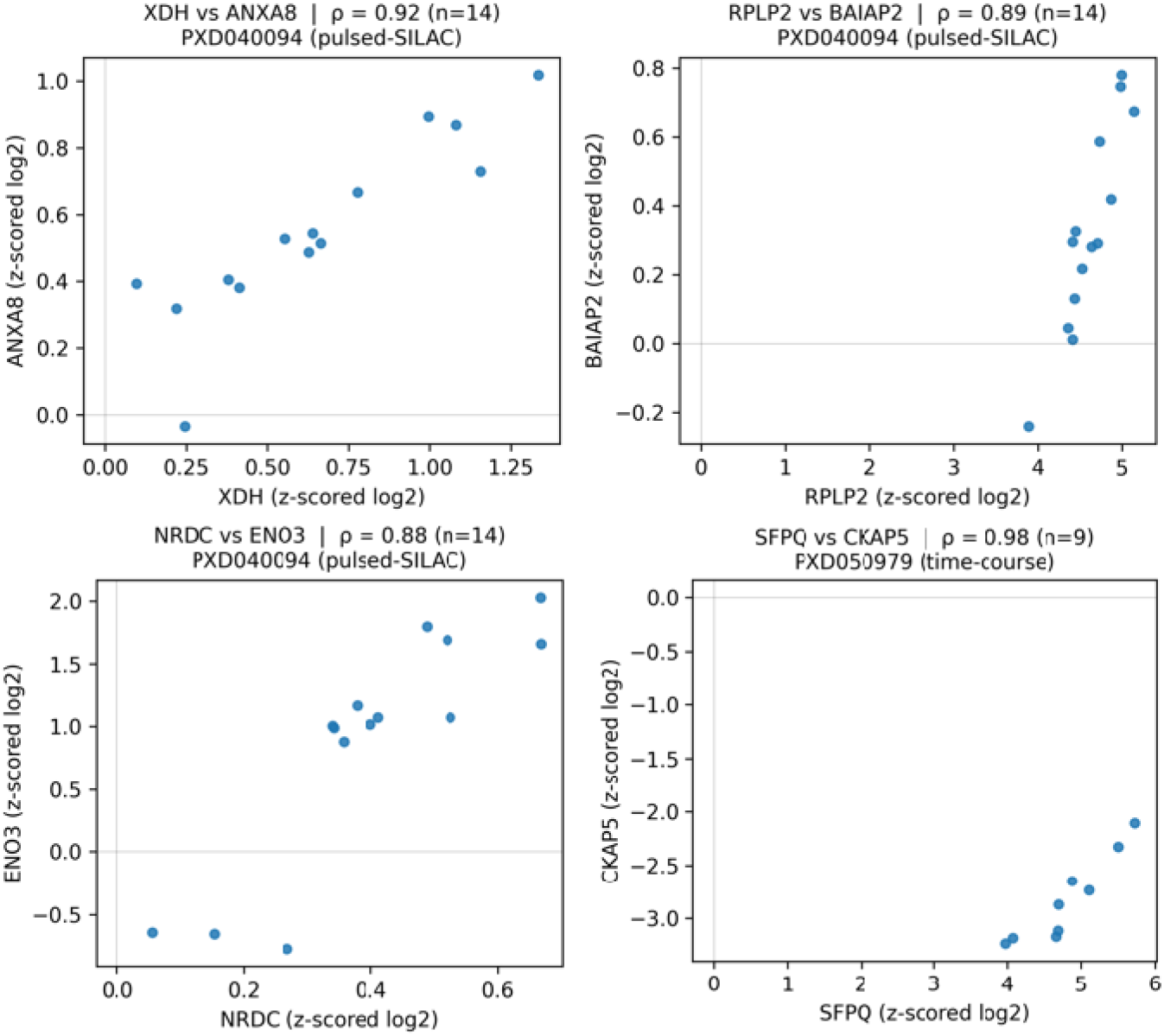
Proteomics support for LazyNet edges in GPX4 loss-of-function fibroblasts. Protein– protein abundance scatterplots for representative LazyNet edges across SILAC channels. Axes show z-scored log_2_ protein levels; each point is one channel/time point. Panels report Spearman ρ and sample count (n). Three pairs come from the pulsed-SILAC GPX4-KO dataset PXD040094— XDH–ANXA8 (ρ = 0.92, n = 14), RPLP2–BAIAP2 (ρ = 0.89, n = 14), NRDC–ENO3 (ρ = 0.88, n = 14) —and one from the tamoxifen-inducible GPX4-KO time course PXD050979—SFPQ–CKAP5 (ρ = 0.98, n = 9). Edges were selected for adequate coverage (≥ 8 quantified channels when available) and clear co-variation.

### 3.6 Biological Insights

The 32×4 subgraph seeded on the CRISPR A/I screen recapitulates the ferroptosis core: among the top 1,000 LazyNet edges (Supplement S1C), 15 of the 27 benchmark regulators reported by Tian et al. cluster near the network center, consistent with the canonical lipid-peroxidation axis [33].

Several connections align with established mechanisms, supporting the face validity of the wiring. A mitochondrial module links MFN2-mediated mitophagy, which constrains ROS-driven death, with SIRT3 signaling that modulates ferroptosis through GPX4 and AMPK–mTOR pathways [37– 38]. Immune–redox crosstalk involving TLR4–STAT3 that amplifies mitochondrial ROS is also recovered [44]. In parallel, ADCY10-driven mitochondrial cAMP has been shown to protect neurons under oxidative stress, matching the directionality of edges observed here [45]. These concordances indicate that LazyNet recovers known biology across mitochondrial, immune, and metabolic axes.

Beyond literature-supported modules, the subgraph nominates cross-axis links that, to our knowledge, are not reported as direct connections and are presented as testable hypotheses. First, a lysosomal–autophagy route from PSAP to mTORC1 (MTOR/RPTOR/RHEB), potentially via ULK1-associated scaffolding (with PPP4R1 as a putative bridge), is consistent with PSAP-dependent lipofuscin accumulation and neuron-specific ferroptosis. Second, high-elasticity edges suggest immune inputs from TLR6 and CCR10 into autophagy/mTOR nodes (ATG13, MTOR). Third, a metabolic coupling is predicted in which TRPM6 (Mg^2+^ channel) and ADCY10 interface with the SIRT3–mTORC1 axis.

Taken together, the results point to an integrated immune–mitochondrial–lysosomal circuit in which mitochondrial quality control (MFN2/SIRT3), innate immune signaling (TLRs), and metabolic/cAMP cues (ADCY10/TRPM6) converge on mTOR-linked autophagy to tune the ferroptotic threshold. Edge-level attributes are provided in Supplement S1C to facilitate replication and follow-up.

## 4. Discussion

This study set out to answer a practical question for small labs running CRISPR A/I screens: can we predict post-perturbation expression accurately, on ordinary hardware, and still learn mechanism? Across two datasets, LazyNet achieves strong predictive ranking (ROC-/PR-AUC) under tight compute, with competitive absolute error and clear generalization to a distinct T-cell slice. The neuronal dataset remains our focus; the T-cell results serve as a sanity check that the approach is not dataset-specific. Relative to transformer and state-space baselines, the gains concentrate in threshold-free ranking while ceding some ground on RMSE and Pearson Correlation in certain settings—an expected outcome when a single per-gene operating point is fixed. Taken together, the results indicate that an interpretable, CPU-tractable ODE update can be competitive with modern sequence models for single-snapshot perturbation prediction.

A central feature of LazyNet is that its one-step, log–exp residual update exposes elasticities— local, directed sensitivities of genes to one another around a baseline. Averaging Jacobians across replicas produces a consensus, directed interaction matrix that we expand into a compact subgraph for interpretation. This graph recovers a substantial fraction of known ferroptosis regulators and shows coherent support across independent resources: enrichment in large-scale co-expression (ARCHS4), non-random overlap with curated functional associations (STRING), and concordant protein-level co-variation in GPX4 loss-of-function proteomes. The agreement is moderate rather than exhaustive — as expected given that elasticity (local, directional quantitative response) and database edges (heterogeneous, often undirected evidence) measure different facets of biology—but it is sufficient to anchor new, testable hypotheses, including a lysosomal–mitochondrial–immune module that nominates specific links into mTOR-connected autophagy.

Equally important is practicality. LazyNet ‘ s capacity is concentrated into a small set of mechanistically meaningful weights, which helps it converge quickly on CPUs without external priors, while still producing parameters that carry direct interpretive value (rate-like elasticities). The ensemble Jacobian pipeline is deterministic and light enough to run end-to-end on a single node, and the entire workflow—preprocessing, splits, commands, and seeds—is packaged for exact replication. In contrast, graph-inference tools optimized for edge ranking from co-expression can generate high-accuracy local subgraphs but are not designed to provide a globally consistent, directed dynamic map aligned with potential causality [55]; for comparative purposes we therefore emphasize module-level enrichment against authoritative, curated resources and orthogonal experiments rather than raw edge-overlap counts.

Several limitations follow from design choices. Working in log space requires positive inputs and introduces mild dependence on normalization and pseudocounts. A single explicit Euler step matches two-snapshot designs but leaves the absolute Δt unidentified and cannot represent delays or oscillations; elasticities are local to the chosen baseline and may vary across states. Our enrichment tests rely on incomplete references whose coverage and evidence channels differ from the signal inferred by LazyNet; consequently, we do not expect high global concordance, and we treat overlaps as supportive rather than definitive. Finally, within a fixed wall-time budget, excessively large models can undertrain and reduce accuracy despite higher nominal capacity.

Looking ahead, three directions appear most impactful. First, time-resolved perturbation series would allow multi-step fitting that estimates Δt, tests stability, and distinguishes fast from slow modes. Second, structured priors (e.g., motif-based regulator sets or pathway constraints) could be added in a transparent way — regularizing the elasticity matrix without sacrificing interpretability. Third, prospective validation—targeted CRISPR edits chosen from LazyNet’s high-elasticity, database-unsupported edges—will be essential to calibrate precision and to refine the expansion rules that balance depth with tractability. Within these bounds, our results suggest that a one-step neural-ODE view can make routine Perturb-seq datasets actionable on commodity hardware, linking accurate prediction with mechanism-level hypotheses that are ready for bench testing.

## 5. Conclusions

Perturbation datasets remain costly to generate at scale and hard to harmonize across studies, so many groups benchmark on public resources while analyzing their own data with bespoke code. Our goal was a method that operates directly on local measurements, runs under modest compute, and yields parameters with clear mechanistic meaning. LazyNet meets this brief: a one-step neural-ODE that trains on CPUs, reports reproducible predictions, and exposes directed elasticities suitable for network interpretation.

Empirically, LazyNet attains competitive or superior predictive accuracy with substantially fewer parameters and lower wall-time than strong baselines, while producing elasticities that show coherent external concordance at the module level. A central advantage is the log–linear–exp map, which exactly represents multiplicative terms. Synergistic, multi-locus effects therefore appear as explicit components rather than opaque composites, enabling practical design and prioritization of combinatorial perturbations from sparsely sampled CRISPR A/I data.

Limits follow from design choices: two-snapshot learning cannot identify absolute Δt, delays, or oscillations; log-domain preprocessing (pseudocount, normalization) introduces mild dependencies; the update is Markovian without latent memory; and full transcriptome-scale expansion can be memory-intensive (partially mitigated with block-sparse top-k storage and streaming). Nonetheless, within a fixed 1-hour budget, the model’s capacity-to-convergence trade-off is favorable on CPUs, and its parameters remain directly interpretable.

Future work will extend validation to time-resolved perturbation series and additional cell contexts, incorporate lightweight structured priors (motifs, pathways) and uncertainty quantification, and perform prospective tests of synergy-driven predictions. Within these bounds, LazyNet offers a reproducible, resource-efficient route to synergy-aware mechanistic inference from perturbation data—bridging accurate prediction with directed, testable hypotheses.

## Supplementary Materials

Supplementary Table S1A — Ferroptosis seed selection. List of seed genes with selection criteria, source references/databases, and brief inclusion rationale (gene symbol, alias, source, evidence note).

Supplementary Table S1B — Subgraph metrics. Global metrics for each expanded subgraph (e.g., BFS 32 × 4): nodes, edges, density, average degree, clustering coefficient, diameter/90th-percentile path length, hub scores, and community counts.

Supplementary Table S1C — Selected subgraph details. Node- and edge-level attributes for the chosen subgraph: gene symbol, aliases, degree, community label; edge elasticity/sign, rank/percentile, and any external matches (ARCHS4 r, STRING score).

Supplementary Table S1D — Per-seed recall. Recall of benchmark ferroptosis regulators per seed (and aggregated), with optional precision@K and top-K per hop settings.

Supplementary Figure S2A — ARCHS4 comparison histogram. Histogram of Pearson’s r for model-inferred edges vs. degree-/expression-matched null pairs; bin width and sample sizes indicated.

Supplementary Table S2B — ARCHS4 comparison statistics. Summary statistics (mean, median, IQR, upper-tail mass ≥r_0_), test results (e.g., Mann–Whitney U / KS), effect sizes (e.g., Cliff’s δ), and adjusted p-values.

Supplementary Table S2C — Pair enrichment. Enrichment of high co-expression pairs at thresholds (e.g., r≥0.1/0.2/0.3), reported as observed/expected, fold-enrichment, 95% CIs, and matched-null definition.

Supplementary Table S3A — STRING comparison overlap. Overlap between inferred edges and STRING v12 interactions across combined-score cutoffs, with counts, hypergeometric p-values, FDR, and evidence-channel breakdown (experimental, database, co-expression, text mining). Supplementary Table S3B — STRING literature support. Curated literature for overlapping edges (gene A–gene B, PMID(s), evidence type/brief note, STRING channel), suitable for spot-checking biological plausibility.

Supplement S4 — Mathematical Details of LazyNet. Formal derivation of LazyNet’s one-step log –exp residual update; exact representation of multiplicative synergies.

## Abbreviations

The following abbreviations are used in this manuscript:

A/I: CRISPR activation/interference
AIVC: Activation/Interference variable combinations (multi-perturbation synergy)
ARCHS4: Large-scale RNA-seq co-expression resource
BFS: Breadth-first search
CRISPR: Clustered Regularly Interspaced Short Palindromic Repeats
GPX4: Glutathione peroxidase 4
KO: Knockout
LLE: Log–Linear–Exp map (LazyNet block)
MAE: Mean absolute error
ODE: Ordinary differential equation
r: Pearson correlation coefficient
ResNet: Residual network
RMSE: Root mean square error
scRNA-seq: Single-cell RNA sequencing
SILAC: Stable isotope labeling by amino acids in cell culture
STRING: Search Tool for the Retrieval of Interacting Genes/Proteins
HVG: Highly variable genes

