## supplement_v2 for "LazyNet: Interpretable ODE Modeling of Sparse CRISPR Single-Cell Screens Reveals New Biological Insights": Supplement_S4.docx

Supplement S4 — Mathematical Details of LazyNet and Elasticity‑Based Network Inference

**S4.1 Notation and data model**

Let x ∈ ℝ>0^p denote gene expression (counts or normalized abundances), u ∈ {0,1}^p an intervention indicator (one‑hot or multi‑hot guide vector), and ε>0 a small pseudocount ensuring positivity. We work in the log domain so that fold‑changes become additive. All logarithms are elementwise.

z = log(x + ε) ∈ ℝ^p	(S4.1)

For two‑snapshot experiments, each cell provides a triple (z_pre, u, z_post); “pre”/“post” refer to the baseline and post‑perturbation states after library‑size normalization and pseudocount shift.

**S4.2 One‑step log–exp residual update (Euler discretization)**

LazyNet instantiates a single explicit ODE step in the log domain via a residual map gθ(·):

z_post = z_pre + gθ(z_pre, u)	(S4.2)

In expression space, the residual is multiplicative due to the log–exp change of variables:

x_post = (x_pre + ε) ⊙ exp( gθ( log(x_pre + ε), u ) ) − ε	(S4.3)

Equation (S4.2) is the Euler step of ẋ in log space with an unknown step size Δt. Writing fθ for the continuous‑time field,

dz/dt = fθ(z, u), gθ(z, u) = Δt · fθ(z, u)	(S4.4)

Hence only the product Δt·fθ is identifiable from two snapshots; absolute Δt is not determined.

**S4.3 Exact representation of multiplicative synergy via the log–linear–exp map**

If the residual is affine in (z, u), gθ(z, u) = A z + B u + b, then a linear form in z produces exact power‑law interactions in x (elementwise):

x_post,i = (x_pre,i + ε) · Π_j (x_pre,j + ε)^{A_{ij}} · exp( Σ_k B_{ik} u_k + b_i ) − ε	(S4.5)

Thus pairwise and higher‑order multiplicative effects appear as explicit model components. With smooth hidden layers, gθ represents mixtures of power laws (log‑sum‑exp features), preserving multiplicative semantics.

**S4.4 Training objective (robust regression)**

Main runs use the Huber loss with δ = 0.1 on the modeling gene panel 𝒢. For residual r = z_post − z_pre − gθ(z_pre,u):

L_Huber(θ) = (1/|𝔅||𝒢|) · Σ_{(·)∈𝔅} Σ_{i∈𝒢} [ ½ r_i^2 if |r_i|≤δ ; δ(|r_i| − ½δ) otherwise ]	(S4.6)

The Huber objective soft‑clips rare, extreme deviations common in perturbation screens while remaining quadratic in the bulk.

**S4.5 Elasticities and Jacobians (directed, mechanism‑level edges)**

Define the log‑elasticity matrix at a baseline b (a fixed z‑vector, e.g., a cohort mean):

E(b,u) := ∂z_post/∂z_pre |_{(z_pre=b,u)} = I + J(b,u), with	J(b,u) := ∂gθ/∂z (b,u)	(S4.7)

Entry J_{ij} quantifies the directed local sensitivity of gene i to gene j in log space (holding u fixed). In expression space, ∂ log x'_i / ∂ log x_j ≈ E_{ij}(b,u). We rank |J_{ij}| to summarize influence magnitudes.

**S4.6 Ensemble Jacobians and subgraph extraction**

Given R independently seeded checkpoints {θ^(r)}_{r=1..R}, compute J^(r)=∂gθ^(r)/∂z at baseline b (autodiff). The ensemble Jacobian is the elementwise average:

J̄ = (1/R) · Σ_{r=1}^R J^(r)	(S4.8)

Because differentiation and expectation commute under mild conditions, J̄ estimates the expected local sensitivity. For each seed gene s, we rank both downstream (J̄_{·s}) and upstream (J̄_{s·}) magnitudes, retain neighbors with |J|≥10⁻³ and above the 95th or 99th percentile of the ranked list, then grow a breadth‑first 32×4 consensus subgraph. For single‑baseline Jacobians that yield few candidates, we omit additional FDR filtering (q=1) and report the full ranked sets.

**S4.7 Identifiability and invariances**

Time‑step confounding: two snapshots identify Δt·fθ, not Δt itself. Scaling and shifts in x (library size, ε) enter additively in z; the residual learns on the chosen normalization. Elasticities are state‑local (evaluated at b) and may vary with the baseline—an expected property of dynamic models.

**S4.8 Computation and memory**

Forward/backward scale with parameter count. Jacobian columns are obtained via vector‑Jacobian products (autodiff) to avoid forming the full p×p matrix in memory. For storage and ranking, we keep block‑sparse top‑k entries per row/column and stream the remainder. This enables transcriptome‑scale evaluation on CPUs.

**S4.9 Evaluation linkage (module‑level enrichment)**

Curated databases (e.g., STRING, ARCHS4) aggregate heterogeneous and often undirected evidence, so strict edge‑wise overlap is an imperfect target for directed, local elasticities. We therefore evaluate at the module level (pathways/regulons), testing whether the inferred subgraph induces coherent activity/enrichment that aligns with known biology, and we corroborate with orthogonal proteomics when available (e.g., GPX4‑knockout SILAC). This yields a more stable and functionally interpretable assessment than raw edge counts.

**S4.10 Two‑gene toy example (exact multiplicative synergy)**

Let p=2, gθ(z,u)=A z with A = [[0, α],[β, 0]]. Then (S4.5) gives:

x'_1 = (x_1+ε) · (x_2+ε)^α − ε, x'_2 = (x_2+ε) · (x_1+ε)^β − ε	(S4.9)

Even with a linear residual in z, the update in x is multiplicative, explicitly encoding synergy between genes 1 and 2. Local elasticities at baseline b are J = A, so |α| and |β| appear directly in the Jacobian and in the ranked neighbor lists.
