## Supplementary figures and images for "LazyNet: Interpretable ODE Modeling of Sparse CRISPR Single-Cell Screens Reveals New Biological Insights"

### Supplement S2A.png

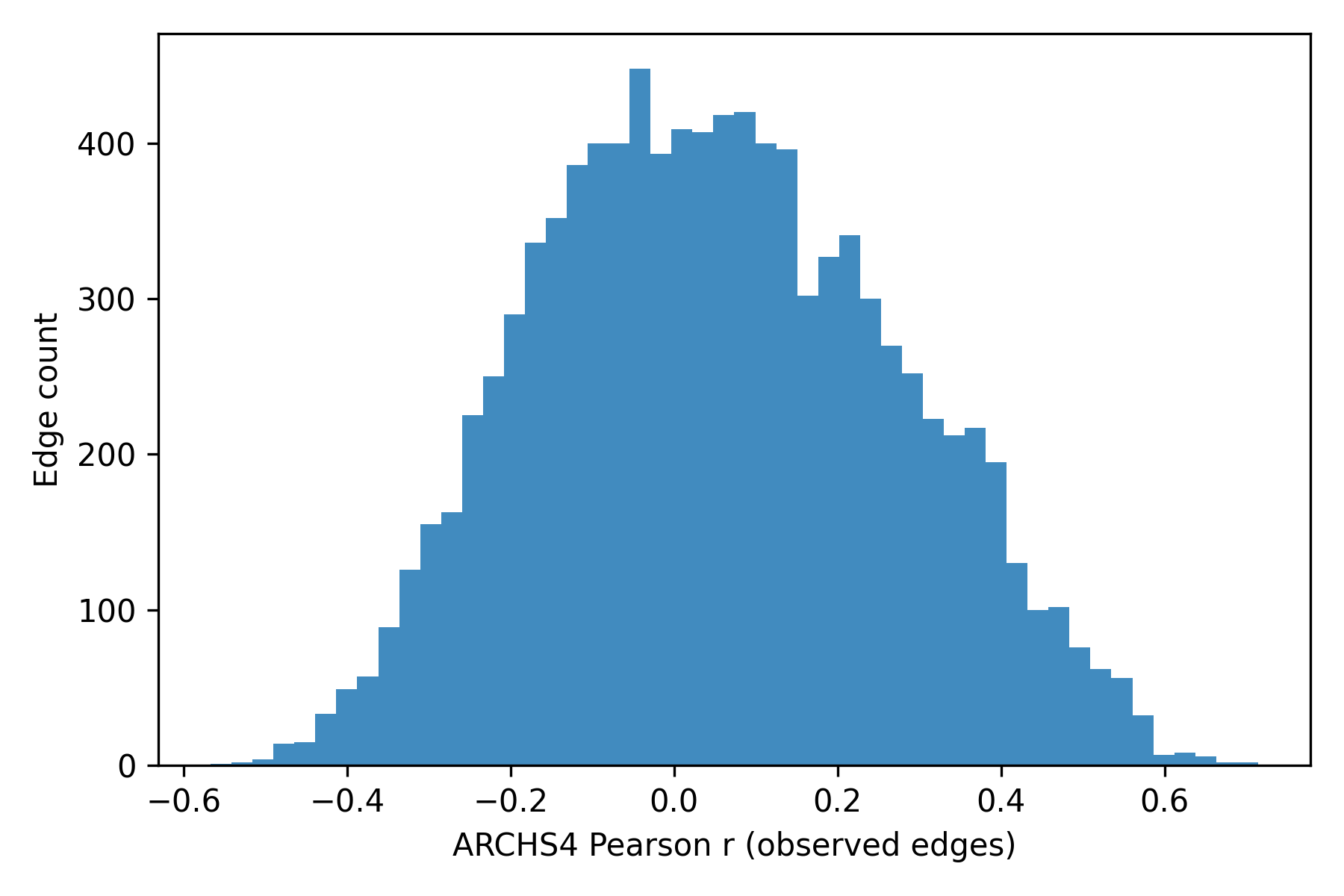
